# High Incidence of Estrous Cycle Irregularities in Heterogeneous Stock (HS) Rats is Associated with Severe Cocaine Addiction-like Behaviors

**DOI:** 10.1101/2025.03.06.641918

**Authors:** Elizabeth A. Sneddon, Supakorn Chonwattanagul, Kathleen Bai, Pranav H. Kurup, Sonja L. Plasil, Michelle R. Doyle, Sélène Zahedi, Sierra Simpson, Benjamin C. Sichel, Dyar N. Othman, Molly Brennan, Abraham A. Palmer, Marsida Kallupi, Lieselot L.G. Carrette, Giordano de Guglielmo, Olivier George

**Affiliations:** Department of Psychiatry, University of California San Diego, La Jolla CA 92093, USA; The Scripps Research Institute, La Jolla, CA 92037; Institute for Genomic Medicine, University of California San Diego, La Jolla CA 92093, USA; Institut de Neurosciences de la Timone, Aix-Marseille Université, Marseille, 13005, France

## Abstract

Hormonal fluctuations throughout the estrous cycle have been hypothesized to influence drug-related behaviors. Preclinical models show that some cocaine-related behaviors are influenced by the estrous cycle. However, the extent to which the estrous cycle modulates cocaine self-administration in outbred heterogeneous stock (HS) rats, a population that captures human genetic diversity, is unknown. This study aimed to examine the relationship between estrous phases and cocaine self-administration behavior in HS rats using a model of extended access to cocaine self-administration. We focused on the escalation of intake, breaking point, and resistance to foot shock. Using vaginal swabbing and lavage techniques, we first characterized the relationship between estrous phase and cocaine intake. We then comprehensively evaluated estrous cycling patterns in young adult and adult HS rats, comparing them with Wistar rats. Contrary to our hypothesis, estrous phase showed no association with cocaine self-administration in HS rats. HS rats exhibited irregular estrous cycling with variability to the phase length, even in the absence of drug exposure, a phenomenon not observed in the Wistar strain. Irregular estrous cycle was associated with high cocaine-related behaviors. This study provides the first evidence that some female HS rats exhibit irregular estrous cycling. Moreover, rats with severe addiction-like behaviors had more instances of irregular cycling. These results demonstrate that, in HS rats, the estrous phase *per se* has no major influence on cocaine self-administration, but that the severity of addiction-like behaviors are associated with more irregularity of the estrus cycle. As HS rats gain popularity in behavioral and genome-wide studies, understanding these cycle disruptions is crucial as they may reveal genetic links into female vulnerability to drugs.

## Introduction

In the United States, 32.1 million women have substance use or mental health disorders [1]. Over the past decade, the rate of cocaine use has increased steadily for women, who more rapidly progress from initial use to dependence than men [2]. Women report stronger pleasurable effects [3] and craving during abstinence [4] compared to men. However, the mechanisms for these differences are unclear.

Hormonal fluctuations across the menstrual cycle (∼28 days) [5] may influence drug-related behaviors. While some studies report no differences of smoked cocaine use between the follicular (high estrogen) and luteal phases [6], women in the follicular phase rate cocaine as more pleasurable compared to those in the luteal phase [6]. Cravings from cocaine- or stress-related cues are also stronger during the follicular phase [7–9]. Intranasal cocaine results in higher plasma cocaine levels during the follicular phase [10], though this is not observed for intravenous cocaine [11]. These findings suggest hormonal fluctuations may play a critical role in cocaine use disorder in women.

Rodents show a similar pattern of fluctuating hormone levels in their estrous cycle (4-5 days), which consists of proestrus, estrus, metestrus, and diestrus [12]. Self-administration varies during the estrous cycle, where rats show greater motivation and increased cocaine-seeking during the estrus phase [13–17]. However, not all studies support this; one found no effect of estrous phase on preference for cocaine versus food [18]. Additionally, disruptions to the normal estrous cycle have been documented following cocaine self-administration in rodents [13,19–21]. Despite these associations, the relationship between estrous phases and cocaine-related behaviors needs to be better established.

Heterogeneous stock (HS) rats are valuable for investigating the genetic variation underlying drug-related behaviors. We and others have characterized a range of addiction-related behaviors for both drug (e.g., cocaine and oxycodone) and non-drug (e.g., food) rewards [22–29]. Sex differences in self-administration for both cocaine [24] and oxycodone [26] have been identified in HS rats. Despite the strength of this model, it has yet to be validated if the estrous cycle is consistent with other strains and if specific phases are associated with cocaine-related behaviors.

We aimed to characterize the estrous cycle in HS rats and assess its association with cocaine-related behaviors. Vaginal samples or lavages were collected at multiple time points across experiments. In Experiment 1, rats underwent cocaine self-administration, with samples collected before and during drug exposure. Experiment 2 further examined estrous cycling in young and adult HS rats at various timepoints over five consecutive days. To validate our methods, Experiment 3 assessed estrous cycling in adult Wistar rats with a similar sampling protocol. We hypothesized that HS rats would show increased cocaine use during high-estrogen phases (estrus + proestrus). This study aimed to clarify how hormonal fluctuations influence cocaine-related behaviors in this translational model. Here, we show that female HS rats with irregular estrous cycling may influence cocaine vulnerability, even in the absence of direct phase associations on single swabbing days.

## Methods

Full procedures are detailed in the George lab protocol repository: https://www.protocols.io/workspaces/george-lab.

### Subjects

411 female HS rats were obtained from Wake Forest University (WFU; n = 256) and UC San Diego (n = 155). Ten female Wistar rats were obtained from Charles River for control experiments. HS rats shipped from WFU arrived at UC San Diego at 3–4 weeks of age and Wistars at 6 weeks. Upon arrival from WFU, rats underwent a 2-week quarantine, then were housed in pairs under a 12-hour light/dark cycle at controlled temperature (20–22°C) and humidity (45–55%). Rats had *ad libitum* access to tap water and food (PJ Noyes Company, Lancaster, NH, USA) and were handled for at least five days before experiments. All procedures followed NIH guidelines and were approved by the UC San Diego IACUC committee.

### Estrous cycle monitoring

For Experiment 1, HS rats (n = 306) were swabbed using a sterile cotton swab dipped in Milli-Q® water one hour before drug exposure, either on the final short access (ShA 10) or the first progressive ratio (PR1) session, and the last long access (LgA 14) (**Fig 1A**). The vaginal cells were collected and placed on a glass microscope slide. In a smaller cohort of 39 HS rats, 28 of which received cocaine while 11 remained cocaine-naive, vaginal samples were collected for four consecutive days every 24 hours before the drug self-administration paradigm began.

**Figure 1.**
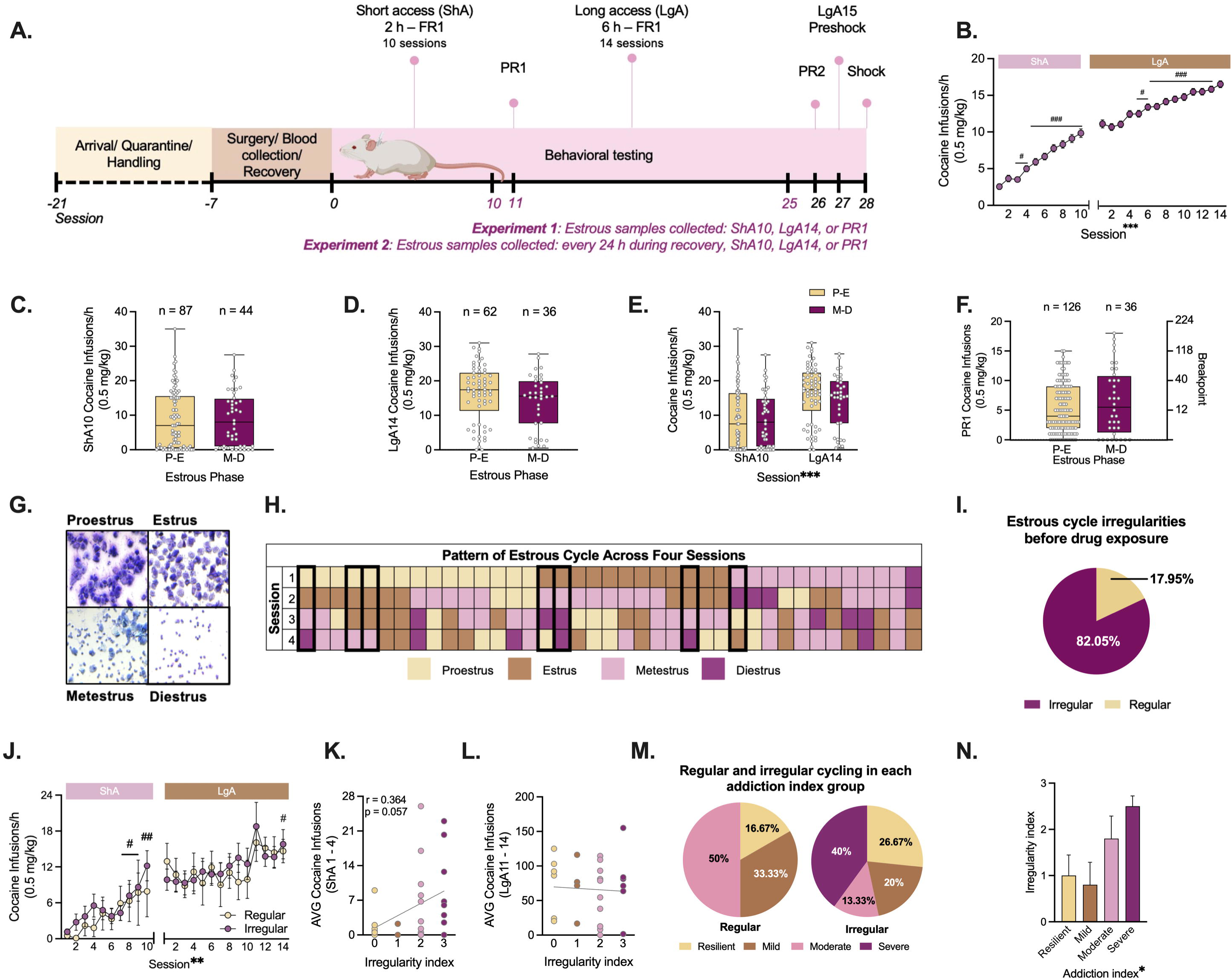
Irregular estrous cycling may influence cocaine vulnerability. **(A)** Timeline of Experiment 1. **(B)** Timeline of cocaine self-administration (n = 306). **(C)** Estrous cycle phase distribution during short access (ShA) 10 (n = 131). **(D)** Estrous cycle phase distribution during long access (LgA) 14 (n = 98). **(E)** Regardless of the collapsed estrous phase, subjects earned more cocaine infusions during LgA vs. ShA sessions. **(F)** The collapsed estrous phases are not associated with cocaine infusions during progressive ratio 1 (PR1, n = 162). **(G)** Visual representation of cytology in each phase of the estrous cycle: diestrus (D), proestrus (P), estrus (E), and metestrus (M). **(H)** Visualization of 24-h cycling in a subset of HS rats (n = 39). Regular cycling is outlined by a bolded line. **(I)** Percentage of rats with regular vs. irregular cycling. **(J)** Self-administration of rats showing regular (n = 6) and irregular (n = 14) cycling patterns across ShA and LgA sessions. **(K)** Correlation between irregularity index and average infusions on ShA sessions 1-4 (p = 0.057). **(L)** No correlation between the irregularity index and cocaine infusions during the last four sessions of LgA. **(M)** Percentage of rats with regular (left) or irregular (right) cycling based on their addiction index. **(N)** Distribution of rats by the irregularity and addiction indices. (* p < 0.05, ** p < 0.001, or **** p < 0.001, main effect One- or Two-Way ANOVA. # p< 0.05, ## p < 0.001, or ### p < 0.001, session compared to session 1, Dunnett’s).

For Experiment 2, the estrous cycle was monitored in sexually mature [30] female HS rats (n = 81) at 7-8 weeks of age. Samples were taken every 7-9 hours across 5 days (**Fig. 2A**). As the multiple time points of samples were being collected, vaginal lavage was used since this method is less invasive [31,32], and has shown consistent findings in past work [33]. An additional cohort of rats (n = 24) received vaginal lavage at 10-11 weeks of age to assess if the estrous cycle stabilized in adulthood.

**Figure 2.**
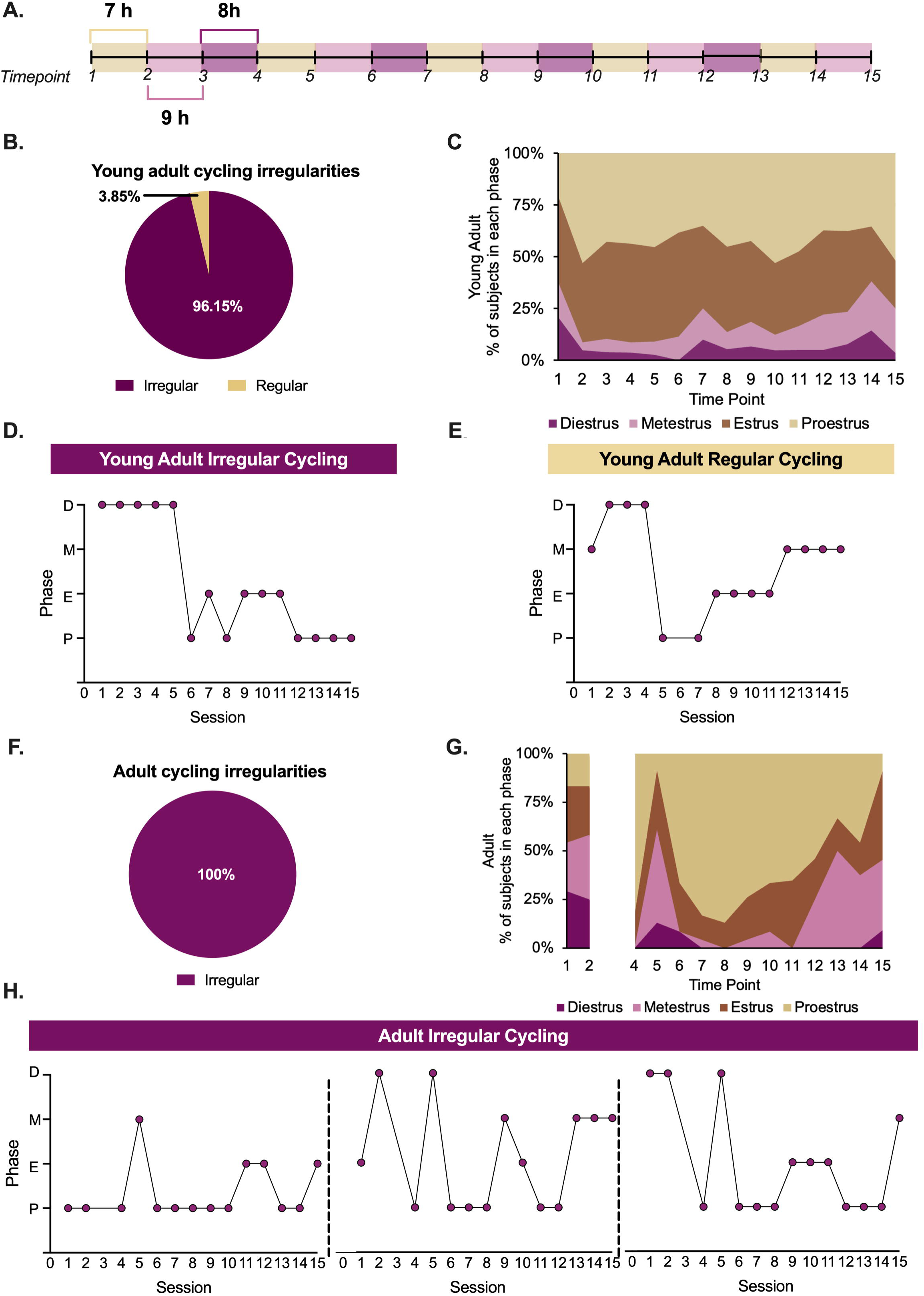
Young adult and adult female HS rats show irregular estrous cycling without drug exposure. **(A)** Timeline of Experiments 2, sampling of young adult (7 – 8 weeks, n = 81) and adult (10 – 11, n = 24). **(B)** Percentage of rats with regular vs. irregular cycling in young adults. **(C)** Percentage of young adults in each phase across time points. Representative line graphs showing **(D)** irregular and **(E)** regular cycling in young adult HS rats. **(F)** Percentage of rats with irregular cycling in adults. **(G)** Percentage of adults in each phase across time points. **(H)** Representative line graphs showing irregular cycle in adults.

For Experiment 3, female Wistar rats (n = 10) vaginal lavage was used for 5 days every 12 hours as a control (**Fig. 3A**). This schedule of sampling and strain was used as Wistars have been validated as having consistent estrous cycling with up to 60-70% of rats showing regular estrous cycle [34].

Unstained slides (n = 306) were visualized in Experiment 1. All other samples were stained 24 hours post-collection using a Hema 3 stat pack (Fisher Scientific, Pittsburgh, PA), with slides dipped in Fixative (30 sec), Solution I (30 sec), and Solution II (15 sec). Estrous phases were identified using a BZ-X800 Analyzer (Keyence, Itasca, IL) based on cell type distribution [12,33,35,36]. Regular cycles followed established phase durations [12], while irregular cycles exceeded these durations or deviated from typical patterns [34].

### Apparatus & Behavioral Testing

Cocaine self-administration was conducted in operant conditioning chambers (Med Associates, St. Albans, VT, USA) housed in soundproof, ventilated cubicles, as previously described [27–30,32]. Each chamber contained two retractable levers, with a cue light above the active lever, and a floor made of metal rods. Foot shocks (0.3 mA, 0.5 seconds) were delivered through an aversive stimulator (ENV-414S, Med Associates).

Rats underwent 10 short-access (ShA; 2-hour) sessions followed by 14 long-access (LgA; 6-hour) sessions, conducted on weekdays, beginning two hours into the dark cycle, as previously described [24]. Cocaine (0.5 mg/kg/infusion) was delivered intravenously on a fixed ratio schedule, with each infusion followed by a 20-second timeout. Inactive lever responses were recorded. Following self-administration, rats completed progressive ratio (PR) testing after ShA 10 (PR1) and LgA 14 (PR2), where the breakpoint was defined as the last completed ratio in a 60-minute period during which a ratio was not completed. A final one-hour foot shock session was conducted under the fixed ratio conditions, with 30% of cocaine infusions paired with foot shocks.

### Drugs & Surgery

Cocaine HCl (National Institute on Drug Abuse, Bethesda, MD) was dissolved in 0.9% sterile saline. Postoperative care included subcutaneous flunixin (2.5 mg/kg) for analgesia and intramuscular cefazolin (330 mg/kg) for to prevent infection. Catheter patency was maintained with a daily flush of heparin sodium (10 U/mL) and cefazolin in bacteriostatic saline.

Rats were implanted with jugular vein catheters under isoflurane anesthesia (1–5%) using aseptic techniques [27–30,32]. Catheters were inserted into the right jugular vein which was connected to a cannulae secured with dental cement and mesh. The cannulae port was externalized via a dorsal incision on the back of the rodent. Incisions were closed with Vetbond tissue adhesive (Santa Cruz Biotechnology Inc., Dalla, TX), and rats recovered for five days before behavioral testing.

### Data Analysis

For Experiment 1, cocaine infusions were normalized to infusions per hour. Self-administration data were analyzed using repeated measures (RM) One-Way or Two-Way Analysis of Variance (ANOVA) (session or session × group as factors). If no interaction was detected, groups were collapsed for a One-Way ANOVA. In cases of missing values, a Mixed-Effects ANOVA was used. *Post hoc* Dunnett’s tests assessed escalation vs. session 1. Animals that did not escalate in LgA were excluded. To confirm their exclusion did not impact our results, we compared session 1 and 14 between the full (n = 306) and reduced (n = 20) cohorts using an unpaired t-test with Welch’s correction. Greenhouse–Geisser correction was applied when sphericity was violated (ε < 0.75).

Estrous phase effects on ShA, LgA, or PR1 infusions were analyzed using One-Way ANOVA, with phase as a between-subjects factor (all p > 0.05). Given the lack of differences, phases were collapsed into high (proestrus + estrus) vs. low (metestrus + diestrus) estrogen groups and reassessed using an unpaired t-test with Welch’s correction.

To assess the relationship between estrous cycling and cocaine self-administration, an irregularity index was assigned (0–3) based on skipped phases, prolonged durations, or atypical phase sequences [14]. Spearman’s correlation was used to test associations between cycling irregularities and cocaine infusions during early ShA (first four sessions) and late LgA (last four sessions). An addiction index was computed based on quartiles of escalation, motivation, and compulsive-like behavior (averaged Z-scores of three dependent variables explaining ∼50% of the variance) [24]. The distribution of regular vs. irregular cycling across addiction index groups was determined, followed by a One-Way ANOVA to assess the influence of the addiction index.

Estrous phase duration was analyzed using Two-Way ANOVA (estrous phase × timepoint). A subject was considered in a phase if present for at least one timepoint, with cumulative time estimated based on consecutive observations. Phase durations were summarized as mean ± SEM. Percentages of regular vs. irregular cycling subjects were also calculated.

Behavioral data were collected using MED-PC IV software. Imaging data with insufficient samples or contamination (e.g., urine) were excluded. For Experiment 3, data for timepoint 3 were unavailable due to unforeseen circumstances. Two independent researchers verified estrous phase classification. Data analysis and visualization were conducted using GraphPad Prism 10.2.2, RStudio, Microsoft Excel, and BioRender. All values are reported as mean ± SEM unless otherwise stated, with statistical significance set at p < 0.05. Effect sizes (η², 95% CI, and r²) are reported where applicable.

## Results

### Experiment 1: Estrous phase is not associated with cocaine-related behaviors in HS rats

A RM One-Way ANOVA revealed a main effect of session during ShA (F_(4.616, 1366.4)_ = 63.890, p < 0.0001, r^2^ = 0.178). A *post hoc* Dunnett’s test revealed that sessions 2 – 10 were increased compared to session 1 (all p < 0.05). For LgA sessions, a Mixed Effects ANOVA found a main effect of session (F_(6.190, 1831.9)_ = 47.283, p < 0.0001, r^2^ = 0.394). A *post hoc* Dunnett’s test showed that sessions 4 – 14 were higher compared to session 1 (**Fig. 1B**). An unpaired t-test found no differences between phases on the last cocaine ShA session (t_(91.403)_ = 0.085, p = 0.932, r^2^ = 7.974e-005) (**Fig. 1C**) or the last cocaine LgA session (t_(76.093)_ = 1.358, p = 0.178, r^2^ = 0.023) (**Fig. 1D**). When assessing cocaine infusions between the last ShA and LgA session, a Two-Way ANOVA found a significant main effect of session (F_(1, 163)_ = 189.040, p < 0.0001, η^2^ = 0.474) (**Fig. 1E)**. An unpaired t-test found no significant differences between phases on the first PR session (t_(46.838)_ = 1.144, p = 0.258, r^2^ = 0.027) (**Fig. 1F**).

In a subset of these rats (n = 39), we found that 82.05% of the subjects had cycling irregularities and 17.95% exhibited regular cycling (**Fig. 1G-I**). For self-administration sessions, the rats exposed to cocaine were assessed (n = 28), but seven of these were excluded. To ensure this exclusion did not influence our results, we compared sessions 1 and 14 between the larger (n = 306) and smaller (n = 20) cohorts. An unpaired t-test showed no differences between the cohorts for LgA1 (t_(22.854)_ = 0.644, p = 0.526,, r^2^ = 0.018) or LgA14 (t_(20.464)_ = 0.939, p = 0.359,, r^2^ = 0.041). The larger cohort infused 11.20 ± 0.54 of 0.5 mg/kg/hour on LgA1 and 16.62 ± 0.46 of 0.5 mg/kg per hour on LgA14. The smaller cohort infused 10.18 ± 6.84 of 0.5 mg/kg/hour on LgA1 and 14.89 ± 1.78 of 0.5 mg/kg per hour on LgA14. A RM Two-Way ANOVA found a main effect of session for ShA (F_(1.755, 26.329)_ = 6.213, p = 0.008, η^2^ = 0.137) and LgA (F_(3.742, 59.879)_ = 4.116, p = 0.006, η^2^ = 0.006). As no interaction was found, the data were collapsed across groups. A One-Way ANOVA for ShA found a main effect of session (F_(2.838, 45.421)_ = 8.173, p = 0.0002, r^2^ = 0.338). A *post hoc* Dunnett’s test identified that sessions 7 – 10 were escalated compared to session 1 (all p < 0.05). A One-Way ANOVA for LgA found a main effect of sessions (F_(3.795, 64,513)_ = 5.619, p = 0.0007, r^2^ = 0.248). A *post hoc* Dunnett’s test identified that session 14 was escalated compared to session 1 (p < 0.05) (**Fig. J**).

No correlation between irregularity index and cocaine infusions during the first four ShA sessions (r_(26)_ =0.364, p = 0.057, CI: −0.022 to 0.656) (**Fig. 1K**) or the last four LgA sessions was observed (r_(25)_ = −0.137, p = 0.7494, CI: −0.501 to 0.267) (**Fig. 1L**). For rats with regular cycling, 16.67% were characterized as resilient, 33.33% as mild, and 50% as moderate for the addiction index. For the rats with irregular cycling, 26.67% were characterized as resilient, 20% as mild, 13.33% as moderate, and 40% as for the severe addiction index (**Fig. 1M**). When assessing if the addiction index varied by irregularity index, a One-Way ANOVA discovered a main effect of addiction index (F_(3,17)_ = 3.767, p = 0.031, r^2^ = 0.399) (**Fig. 1N**).

### Experiment 2: Estrous cycle irregularities are observed in young adult and adult female HS rats

We found that 96.15% of the young adult rats (n = 78) had cycling irregularities and 3.85% (n = 3) had a regular cycle (**Fig. 2B-E**). A significant main effect of estrous phase was observed when assessing the percentage of subjects in each phase across time point (F_(3, 42)_ = 79.29, p < 0.0001, η^2^ = 84.99) (**Fig. 2C**). When assessing the duration of each estrous phase in young adult female rats, the time spent each phase (in hours) are shown in **Table 1**. When evaluating the percentage of rats that experienced each estrous phase at least once, we observed that 100% of the rats experienced proestrus and estrus, 82.72% experienced metestrus, and 53.09% experienced diestrus.

**Table 1.** Length of Estrous Phases in Young Adult and Adult HS Rats and Adults Wistar Rats (in hours). Average and range of rats were in each phase of the estrous cycle. Data are expressed as mean ± SEM, the median, or ranges. Young adults (n = 81), adult (n = 24), and Wistar (n = 10) rats. (* p < 0.05 vs. Wistars, # p < 0.05 Young Adult vs. Adult HS rats; Holm Sidak’s).

| Estrous phase | Statistical Test | Cycle Length (hours) |  |  |
| --- | --- | --- | --- | --- |
|  |  | Young Adult HS | Adult HS | Adult Wistars |
|  |  | (7 – 8 weeks) | (10 – 11 weeks) | (10 – 11 weeks) |
| Proestrus | *Mean ± SEM* | 18.33 ± 1.19<sup>#</sup> | 27.25 ± 2.59<sup>*</sup> | 12 ± 0 |
|  | *Median* | 17 | 24 | 12 |
|  | *Range* | 7 – 57 | 15 – 56 | 12 – 12 |
| Estrus | *Mean ± SEM* | 19.68 ± 1.40<sup>#</sup> | 10.38 ± 0.78<sup>*</sup> | 22 ± 6.51 |
|  | *Median* | 16 | 9 | 12 |
|  | *Range* | 7 – 63 | 7 – 17 | 12 - 48 |
| Metestrus | *Mean ± SEM* | 9.09 ± 0.36<sup>*</sup> | 7.87 ± 0.35<sup>*</sup> | 16 ± 4 |
|  | *Median* | 8 | 7 | 12 |
|  | *Range* | 7 – 24 | 7 – 15 | 12 – 36 |
| Diestrus | *Mean ± SEM* | 8.71 ± 0.48<sup>*</sup> | 7.57 ± 0.20<sup>*</sup> | 12 ± 0 |
|  | *Median* | 7 | 7 | 12 |
|  | *Range* | 7 – 31 | 7 – 9 | 12 – 12 |

We found that 100% of the adult female rats (n = 24) had cycling irregularities (**Fig. 2F-H**). A significant main effect of estrous phase was observed when assessing the percentage of subjects in each phase across time point (F_(3, 42)_ = 11.05, p < 0.0001, η^2^ = 40.48) (**Fig. 2G**). When assessing the duration of each estrous phase in adult female rats, the time spent in each phase (in hours) are shown in **Table 1**. When evaluating the percentage of rats that experienced each estrous phase at least once, we observed that 100% of the rats experienced proestrus and estrus, while 95.83% experienced metestrus, and 54.17% experienced diestrus.

When comparing the length of the proestrus phase between Wistar, young adult, and adult HS rats, a One-Way ANOVA identified a significant main effect (F_(1, 112)_ = 9.19, p = 0.0002, r^2^ = 0.14). A *post hoc* Holm Sidak’s test revealed that the adult HS rats had a longer proestrus phase compared to the young adult HS rats (p = 0.001) and adult Wistars (p = 0.0008). When assessing the length of the estrus phase between groups, a One-Way ANOVA discovered a significant main effect (F_(1, 112)_ = 5.92, p = 0.0036, r^2^ = 0.10). A *post hoc* Holm Sidak’s test found that the adult HS rats had a shorter estrus compared to the young adult HS rats (p = 0.004) and adult Wistars (p = 0.026). When comparing the length of the metestrus phase between groups, a One-Way ANOVA discovered a significant main effect (F_(1, 112)_ = 11.91, p = 0.004, r^2^ = 0.18). A *post hoc* Holm Sidak’s test identified that young adult (p < 0.0001) and adult (p < 0.0001) HS rats had a shorter metestrus phase compared to adult Wistars. When assessing the length of the diestrus phase between groups, a One-Way ANOVA revealed a significant main effect (F_(1, 112)_ = 5.14, p = 0.007, r^2^ = 0.08). A *post hoc* Holm Sidak’s test identified that young adult (p 0.017) and adult (p = 0.005) HS rats had a shorter diestrus phase compared to adult Wistar rats (**Table 1**).

### Experiment 3: Wistar rats show regular estrous cycling

Sixty percent of the female Wistar rats (n = 10) had regular cycling while the other forty percent did not (**Fig. 3B – D**).

**Figure 4.**
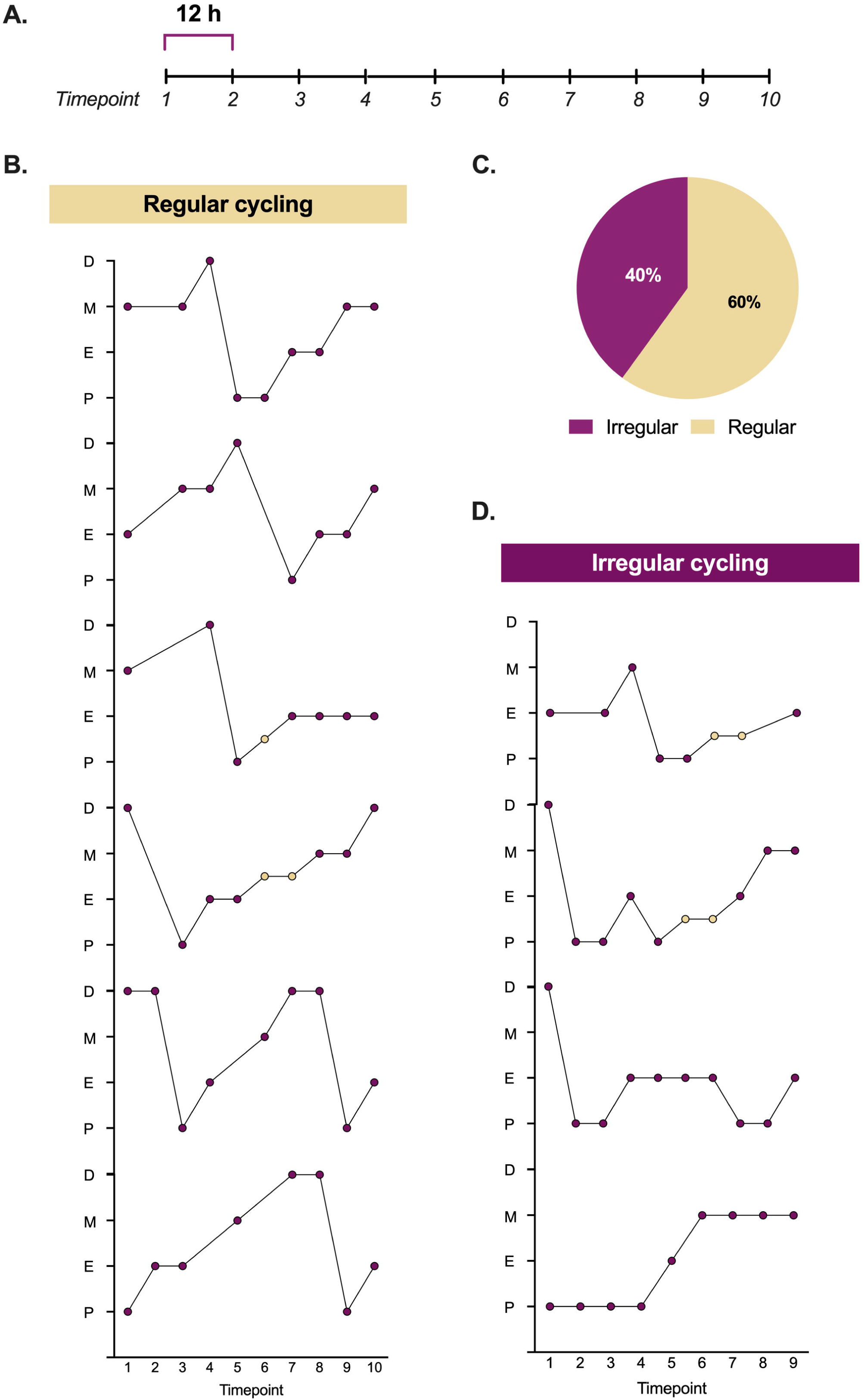
Wistar rats show regular estrous cycling. **(A)** Timeline sample collection. **(B)** Line graph visualization of regular cycling in Wistar rats. **(C)** Percentage of rats with regular vs. irregular cycling. **(D)** Line graph visualization of irregular estrous cycling in Wistar rats. (Tan indicates the transition from one phase to another).

## Discussion

We found that female HS rats with irregular estrous cycling may influence cocaine vulnerability, even in the absence of direct phase associations on single swabbing days. Given the lack of data on estrous cycling in drug-naive HS rats, we conducted additional experiments. HS females lack the regular estrous cycling observed in other outbred lines, such as Wistar rats. These findings conflict with the existing literature and suggest that the HS rats may be a powerful model to investigate estrous cycle irregularities to addiction-related behaviors.

We did not find any associations between estrous phase and cocaine self-administration during ShA10, LgA14, or PR1. These null findings should be interpreted with caution, as a subset of HS rats revealed that only 17.95% of the subjects exhibited regular cycling patterns. We characterized an irregularity index (criteria determined from [34]) to assess whether the irregularities correlated with cocaine-related behaviors. Estrous cycling irregularities before drug exposure nearly met the threshold of significance for self-administration during early short access but showed no association during late long-access sessions. It remains unclear whether cocaine-exposure disrupted the cycle in this study, which may explain the absence of correlations with long access sessions. As irregularities were present before drug exposure, we could not assess if cocaine disrupted the cycle. While some studies link high estradiol to increased cocaine use [14,16,37], others report no association or cue-dependent effects [39,40]. Proestrus and estrus consistently correlate with greater motivation in progressive ratio and reinstatement tests [13,16,38–42]. However, we saw no association with the progressive ratio session, which deviates from these findings.

Since the HS rats are ideal for assessing individual differences [43], we examined whether estrous cycle irregularities correlate with addiction-related behaviors using an addiction index [24,28]. We found that only rats with irregular cycling were classified as severe and estrous irregularities were more common in the vulnerable groups (moderate + severe), suggesting that regular cycling may protect against cocaine pathology. While prior studies show cocaine can disrupt estrous cycling [21,44], this is the first to suggest that pre-existing irregularities may influence later drug intake. Since most studies exclude subjects with irregular cycling [35], our findings offer a unique opportunity to explore cycling irregularities as a potential risk factor for cocaine misuse.

We investigated whether cocaine disrupts the estrous cycling in HS rats. We hypothesized that we would observe regular phase progression with disruptions following cocaine exposure, as previously reported [21,44]. However, we found that 96.15% of young adult and 100% of adult exhibited irregular cycling with no apparent pattern in drug-naïve subjects. The timescale in each phase did not match previous reports of the estrous cycle [12]. We observed that the proestrus and metestrus were extended while estrus and diestrus were blunted in both young adult and adult rats. Proestrus was the most prominent phase, while diestrus was the least, contradicting the literature [12,31,34,45]. Similar proestrus extension has been observed in the Goto-Kakizaki rat strain potentially due to hypothalamic-pituitary-gonadal axis dysregulation [46] which may explain our findings. Our results were surprising as we are unaware of any other rodent model that shows irregularities to this extent and requires further exploration.

Our control experiment with Wistar rats showed 60% regular cycling, consistent with prior reports [34,47]. Given similar irregularity rates in other strains [48–50] with rates between 9 - 40% [34,48,49], methodological error is unlikely. Vaginal swabbing, lavages, and staining were used in our studies in Wistar rats [35] and C57BL/6J mice [33] without any methodological issues.

Although estrous variability is common[51], HS rats show unprecedented irregularities. Cycle length can vary from 3 – 38 days in rats [52,53], though most studies phase across 4 – 5 days [12,31,34]. Cycle and phase length may be different in these rats, but that does not account for cellular variability. Sexual maturation can range from 32 to 34 days to weeks in female rats [54–56]. We initially attributed the irregularities at 7-8 weeks to sexual immaturity or stress from rehousing, but the respective irregularities persisted at 10-11 weeks, suggesting permanence.

While excluding rats with irregular cycling is recommended [32,57] to simplify data collection, this approach does not reflect the human population where 14% - 25% of women experience cycling irregularities [58,59]. Women with menstrual cycle irregularities show 40% higher rates of mental disorders, including substance use disorder [60–71]. As cycling irregularities are associated with mental health outcomes, understanding the factors driving cycling regularity in humans and rodents is crucial. HS rats may be valuable for exploring the genetic factors influencing cycle irregularities and mental health, as we observed that irregular cycling is linked to more severe cocaine addiction-related behaviors. Additionally, HS display increased fear, anxiety, and depressive-like behaviors associated with heightened prolactin levels [72–74], though no estrous cycle associations were measured. A meta-analysis found that increased anxiety-like behaviors are tied to lower estrogen levels [75]. The shortened or skipped metestrus and diestrus phases in female HS rats may lead to increased exposure to estrogen due to the more frequent occurrence of estrus and proestrus phases, potentially affecting their anxiety profile.

These results are based on vaginal cytology, not direct hormonal measurement. While this does limit our conclusions, vaginal cytology remains the most reliable method to assess estrous fluctuations in rodents [32]. Serum hormone measurement was considered, but due to irregular cycling in HS rats, determining an appropriate non-invasive blood collection schedule was not feasible for accurately characterizing the cycle. Additionally, ELISA measurement of estradiol in rodents can vary [76], supporting the continued assessment of vaginal smears. As such, vaginal smears were the most accurate assessment of the estrous cycle at our disposal.

This is the first study to show irregular estrous cycles in female HS rats, potentially contributing to the severity of addiction-like behaviors. These irregularities exist without drug exposure, warranting further genetic and hormonal investigations. As HS rats gain popularity in behavioral and genome-wide studies, understanding these cycle disruptions is crucial as they may reveal genetic links into female vulnerability to drugs.

## Data availability statement

All images and data for this project are available from the corresponding author upon request.

## Acknowledgments

The authors would like to thank the Preclinical Addiction Research Consortium at UCSD.

## Authors’ contributions

Conceptualization: EAS and OG. Methodology: EAS and OG. Formal analysis: ES, SC, SS, and SZ. Investigation: EAS, SC, KB, PK, SLP, MRD, BCS, DNO, and MB. Writing – Original Draft: EAS, SC, and KB. Writing – Review and Editing: EAS, SC, KB, SLP, MRD, SZ, SS, GdG, MK, LLGC, AAP, and OG. Visualization: ES and SZ. Supervision: EAS and OG. Resources: AAP. Project Administration: EAS, MB, GdG, MK, LLGC, and OG. Funding acquisition: EAS, LLGC, and OG.

## Competing interests

The authors declare no competing interests.

[TAB]

## References

1. Bustamante J. NCDAS: Substance Abuse and Addiction Statistics [2021]. 2020. https://drugabusestatistics.org/. Accessed November 18, 2021.

2. Kerver HN, Becker JB. Sex Differences in the Effects and Actions of Cocaine. The Neuroscience of Cocaine. 2017:11–19.

3. Becker JB, Hu M. Sex differences in drug abuse. Front Neuroendocrinol. 2008;29:36–47.

4. Elman I, Karlsgodt KH, Gastfriend DR. Gender differences in cocaine craving among non-treatment-seeking individuals with cocaine dependence. Am J Drug Alcohol Abuse. 2001;27:193–202.

5. Mihm M, Gangooly S, Muttukrishna S. The normal menstrual cycle in women. Anim Reprod Sci. 2011;124:229–236.

6. Evans SM, Haney M, Foltin RW. The effects of smoked cocaine during the follicular and luteal phases of the menstrual cycle in women. Psychopharmacology (Berl). 2002;159:397–406.

7. Moran-Santa Maria MM, Flanagan J, Brady K. Ovarian hormones and drug abuse. Curr Psychiatry Rep. 2014;16:511.

8. Sinha R, Fox H, Hong K-I, Sofuoglu M, Morgan PT, Bergquist KT. Sex steroid hormones, stress response, and drug craving in cocaine-dependent women: implications for relapse susceptibility. Exp Clin Psychopharmacol. 2007;15:445–452.

9. Reed BG, Carr BR. The Normal Menstrual Cycle and the Control of Ovulation. In: Feingold KR, Anawalt B, Blackman MR, Boyce A, Chrousos G, Corpas E, et al., editors. Endotext, South Dartmouth (MA): MDText.com, Inc.; 2018.

10. Lukas SE, Sholar M, Lundahl LH, Lamas X, Kouri E, Wines JD, et al. Sex differences in plasma cocaine levels and subjective effects after acute cocaine administration in human volunteers. Psychopharmacology. 1996;125:346–354.

11. Mendelson JH, Mello NK, Sholar MB, Siegel AJ, Kaufman MJ, Levin JM, et al. Cocaine pharmacokinetics in men and in women during the follicular and luteal phases of the menstrual cycle. Neuropsychopharmacology. 1999;21:294–303.

12. Ajayi AF, Akhigbe RE. Staging of the estrous cycle and induction of estrus in experimental rodents: an update. Fertil Res Pract. 2020;6:5.

13. Roberts DC, Bennett SA, Vickers GJ. The estrous cycle affects cocaine self-administration on a progressive ratio schedule in rats. Psychopharmacology. 1989;98:408–411.

14. Lynch WJ, Arizzi MN, Carroll ME. Effects of sex and the estrous cycle on regulation of intravenously self-administered cocaine in rats. Psychopharmacology. 2000;152:132–139.

15. Carroll ME, Morgan AD, Lynch WJ, Campbell UC, Dess NK. Intravenous cocaine and heroin self-administration in rats selectively bred for differential saccharin intake: phenotype and sex differences. Psychopharmacology. 2002;161:304–313.

16. Feltenstein MW, See RE. Plasma progesterone levels and cocaine-seeking in freely cycling female rats across the estrous cycle. Drug Alcohol Depend. 2007;89:183–189.

17. Kohtz AS, Lin B, Davies H, Presker M, Aston-Jones G. Hormonal milieu drives economic demand for cocaine in female rats. Neuropsychopharmacology. 2022;47:1484–1492.

18. Kerstetter KA, Ballis MA, Duffin-Lutgen S, Carr AE, Behrens AM, Kippin TE. Sex differences in selecting between food and cocaine reinforcement are mediated by estrogen. Neuropsychopharmacology. 2012;37:2605–2614.

19. Grimm J. Cocaine Self-Administration in Ovariectomized Rats is Predicted by Response to Novelty, Attenuated by 17-? Estradiol, and Associated With Abnormal Vaginal Cytology. Physiology & Behavior. 1997;61:755–761.

20. Fuchs RA, Evans KA, Mehta RH, Case JM, See RE. Influence of sex and estrous cyclicity on conditioned cue-induced reinstatement of cocaine-seeking behavior in rats. Psychopharmacology. 2005;179:662–672.

21. Truckenbrod LM, Cooper EM, Wheeler A-R, Orsini CA. Cocaine intake correlates with risk-taking behavior and affects estrous cycling in female Sprague-Dawley rats. Front Behav Neurosci. 2023;17:1293226.

22. Woods LCS, Mott R. Heterogeneous Stock populations for analysis of complex traits. Methods Mol Biol. 2017;1488:31–44.

23. King CP, Tripi JA, Hughson AR, Horvath AP, Lamparelli AC, Holl KL, et al. Sensitivity to food and cocaine cues are independent traits in a large sample of heterogeneous stock rats. Sci Rep. 2021;11:2223.

24. de Guglielmo G, Carrette L, Kallupi M, Brennan M, Boomhower B, Maturin L, et al. Large-scale characterization of cocaine addiction-like behaviors reveals that escalation of intake, aversion-resistant responding, and breaking-points are highly correlated measures of the same construct. Elife. 2024;12.

25. Carrette LLG, Corral C, Boomhower B, Brennan M, Crook C, Ortez C, et al. Leptin protects against the development and expression of cocaine addiction-like behavior in heterogeneous stock rats. Front Behav Neurosci. 2022;16:832899.

26. Kallupi M, de Guglielmo G, Carrette LLG, Simpson S, Kononoff J, Kimbrough A, et al. Individual differences in oxycodone addiction-like behaviors in a large cohort of heterogeneous stock (HS) rats. BioRxiv. 2022.

27. Kallupi M, Carrette LLG, Kononoff J, Solberg Woods LC, Palmer AA, Schweitzer P, et al. Nociceptin attenuates the escalation of oxycodone self-administration by normalizing CeA–GABA transmission in highly addicted rats. Proc Natl Acad Sci U S A. 2020;117:2140–2148.

28. Carrette LL, de Guglielmo G, Kallupi M, Maturin L, Brennan M, Boomhower B, et al. The cocaine and oxycodone biobanks, two repositories from genetically diverse and behaviorally characterized rats for the study of addiction. ENeuro. 2021. April 15, 2021. 10.1523/ENEURO.0033-21.2021.

29. Sedighim S, Carrette LL, Venniro M, Shaham Y, de Guglielmo G, George O. Individual differences in addiction-like behaviors and choice between cocaine versus food in Heterogeneous Stock rats. Psychopharmacology (Berl). 2021;238:3423–3433.

30. Sengupta P. The laboratory rat: Relating its age with human’s. Int J Prev Med. 2013;4:624–630.

31. Cora MC, Kooistra L, Travlos G. Vaginal cytology of the laboratory rat and mouse: Review and criteria for the staging of the estrous cycle using stained vaginal smears. Toxicol Pathol. 2015;43:776– 793.

32. Dalla C, Jaric I, Pavlidi P, Hodes GE, Kokras N, Bespalov A, et al. Practical solutions for including sex as a biological variable (SABV) in preclinical neuropsychopharmacological research. J Neurosci Methods. 2024;401:110003.

33. Sneddon EA, Masters BM, Ream KD, Fennell KA, DeMedio JN, Cash MM, et al. Sex chromosome and gonadal hormone contributions to binge-like and aversion-resistant ethanol drinking behaviors in Four Core Genotypes mice. Front Psychiatry. 2023;14:1098387.

34. Marcondes FK, Bianchi FJ, Tanno AP. Determination of the estrous cycle phases of rats: some helpful considerations. Braz J Biol. 2002;62:609–614.

35. Doyle MR, Dirik S, Martinez AR, Hughes TE, Iyer MR, Sneddon EA, et al. Catechol-O-Methyltransferase inhibition and alcohol use disorder: Evaluating the efficacy of tolcapone in ethanol-dependent rats. Neuropharmacology. 2023;242:109770.

36. Waynforth HB, Flecknell PA. Experimental and surgical technique in the rat. 1992;346.

37. Feltenstein MW, Byrd EA, Henderson AR, See RE. Attenuation of cocaine-seeking by progesterone treatment in female rats. Psychoneuroendocrinology. 2009;34:343–352.

38. Lynch WJ. Acquisition and maintenance of cocaine self-administration in adolescent rats: effects of sex and gonadal hormones. Psychopharmacology. 2008;197:237–246.

39. Lacy RT, Strickland JC, Feinstein MA, Robinson AM, Smith MA. The effects of sex, estrous cycle, and social contact on cocaine and heroin self-administration in rats. Psychopharmacology. 2016;233:3201–3210.

40. Doncheck EM, Liddiard GT, Konrath CD, Liu X, Yu L, Urbanik LA, et al. Sex, stress, and prefrontal cortex: influence of biological sex on stress-promoted cocaine seeking. Neuropsychopharmacology. 2020;45:1974–1985.

41. Nicolas C, Russell TI, Pierce AF, Maldera S, Holley A, You Z-B, et al. Incubation of cocaine craving after intermittent-access self-administration: Sex differences and estrous cycle. Biol Psychiatry. 2019;85:915–924.

42. Corbett CM, Dunn E, Loweth JA. Effects of sex and estrous cycle on the time course of incubation of cue-induced craving following extended-access cocaine self-administration. ENeuro. 2021;8:ENEURO.0054-21.2021.

43. Solberg Woods LC, Palmer AA. Using heterogeneous stocks for fine-mapping genetically complex traits. Methods Mol Biol. 2019;2018:233–247.

44. Raap DK, Morin B, Medici CN, Smith RF. Adolescent cocaine and injection stress effects on the estrous cycle. Physiol Behav. 2000;70:417–424.

45. Goldman JM, Murr AS, Cooper RL. The rodent estrous cycle: characterization of vaginal cytology and its utility in toxicological studies. Birth Defects Res B Dev Reprod Toxicol. 2007;80:84–97.

46. Pinto-Souza ARW, Firetto C, Pérez-Arana G, Lechuga-Sancho AM, Prada-Oliveira JA. Differences in the estrous cycles of Goto-Kakizaki and Wistar rats. Lab Anim (NY). 2016;45:143–148.

47. Elsayed DH, Helmy SA, Dessouki AA, El-Nahla AM, Abdelrazek HMA, El-Hak HNG. Influence of genistein and diadizine on regularity of estrous cycle in cyclic female Wistar rat: interaction with estradiol receptors and vascular endothelial growth factor. Open Vet J. 2022;12:639–648.

48. Karim BO, Landolfi JA, Christian A, Ricart-Arbona R, Qiu W, McAlonis M, et al. Estrous cycle and ovarian changes in a rat mammary carcinogenesis model after irradiation, tamoxifen chemoprevention, and aging. Comp Med. 2003;53:532–538.

49. Mourlon V, Naudon L, Giros B, Crumeyrolle-Arias M, Daugé V. Early stress leads to effects on estrous cycle and differential responses to stress. Physiol Behav. 2011;102:304–310.

50. Schuh KM, Ahmed J, Kwak E, Xu CX, Davis TT, Aronoff CB, et al. A mouse model of oral contraceptive exposure: Depression, motivation, and the stress response. Horm Behav. 2024;158:105470.

51. Robert H, Ferguson L, Reins O, Greco T, Prins ML, Folkerts M. Rodent estrous cycle monitoring utilizing vaginal lavage: No such thing as a normal cycle. J Vis Exp. 2021. August 30, 2021. 10.3791/62884.

52. Long JA, Evans HML. The oestrous cycle in the rat and its associated phenomena. Berkeley, CA: University of California Press; 1922.

53. Westwood FR. The Female Rat Reproductive Cycle: A Practical Histological Guide to Staging. Toxicol Pathol. 2008;36:375–384.

54. Lenschow C, Sigl-Glöckner J, Brecht M. Development of rat female genital cortex and control of female puberty by sexual touch. PLoS Biol. 2017;15:e2001283.

55. Lewis EM, Barnett JF Jr, Freshwater L, Hoberman AM, Christian MS. Sexual maturation data for Crl Sprague-Dawley rats: criteria and confounding factors. Drug Chem Toxicol. 2002;25:437–458.

56. Spear LP. The adolescent brain and age-related behavioral manifestations. Neurosci Biobehav Rev. 2000;24:417–463.

57. Holalagoudar S, Kisielewski S, Martini A, Johnson K, Leoni A-L, Demminger C, et al. Rodent estrous cycle pattern: Harmonizing the cycle evaluation and interpretation. Regul Toxicol Pharmacol. 2024;156:105768.

58. Flickr F us on. How many women are affected by menstrual irregularities? https://www.nichd.nih.gov/. https://www.nichd.nih.gov/health/topics/menstruation/conditioninfo/affected. Accessed October 6, 2024.

59. Nobles J, Cannon L, Wilcox AJ. Menstrual irregularity as a biological limit to early pregnancy awareness. Proc Natl Acad Sci U S A. 2022;119:e2113762118.

60. Reilly TJ, Sagnay de la Bastida VC, Joyce DW, Cullen AE, McGuire P. Exacerbation of psychosis during the perimenstrual phase of the menstrual cycle: Systematic review and meta-analysis. Schizophr Bull. 2020;46:78–90.

61. Ajari EE. Connecting the dots between mental and menstrual health: An exploratory review. J Health Rep Technol. 2021;8.

62. Toffol E, Koponen P, Luoto R, Partonen T. Pubertal timing, menstrual irregularity, and mental health: results of a population-based study. Arch Womens Ment Health. 2014;17:127–135.

63. Yu M, Han K, Nam GE. The association between mental health problems and menstrual cycle irregularity among adolescent Korean girls. J Affect Disord. 2017;210:43–48.

64. Nillni YI, Toufexis DJ, Rohan KJ. Anxiety sensitivity, the menstrual cycle, and panic disorder: a putative neuroendocrine and psychological interaction. Clin Psychol Rev. 2011;31:1183–1191.

65. Nillni YI, Wesselink AK, Hatch EE, Mikkelsen EM, Gradus JL, Rothman KJ, et al. Mental health, psychotropic medication use, and menstrual cycle characteristics. Clin Epidemiol. 2018;10:1073– 1082.

66. Barron ML, Flick LH, Cook CA, Homan SM, Campbell C. Associations between psychiatric disorders and menstrual cycle characteristics. Arch Psychiatr Nurs. 2008;22:254–265.

67. Green SA, Graham BM. Symptom fluctuation over the menstrual cycle in anxiety disorders, PTSD, and OCD: a systematic review. Arch Womens Ment Health. 2022;25:71–85.

68. Gleeson PC, Worsley R, Gavrilidis E, Nathoo S, Ng E, Lee S, et al. Menstrual cycle characteristics in women with persistent schizophrenia. Aust N Z J Psychiatry. 2016;50:481–487.

69. Milano W, Ambrosio P, Carizzone F, De Biasio V, Foia MG, Saetta B, et al. Menstrual disorders related to eating disorders. Endocr Metab Immune Disord Drug Targets. 2022;22:471–480.

70. Poyastro Pinheiro A, Thornton LM, Plotonicov KH, Tozzi F, Klump KL, Berrettini WH, et al. Patterns of menstrual disturbance in eating disorders. Int J Eat Disord. 2007;40:424–434.

71. Algars M, Huang L, Von Holle AF, Peat CM, Thornton LM, Lichtenstein P, et al. Binge eating and menstrual dysfunction. J Psychosom Res. 2014;76:19–22.

72. Díaz-Morán S, Martínez-Membrives E, López-Aumatell R, Cañete T, Blázquez G, Palencia M, et al. What can we learn on rodent fearfulness/anxiety from the genetically heterogeneous NIH-HS rat stock? Open J Psychiatr. 2013;03:238–250.

73. López-Aumatell R, Martínez-Membrives E, Vicens-Costa E, Cañete T, Blázquez G, Mont-Cardona C, et al. Effects of environmental and physiological covariates on sex differences in unconditioned and conditioned anxiety and fear in a large sample of genetically heterogeneous (N/Nih-HS) rats. Behav Brain Funct. 2011;7:48.

74. Lopez-Aumatell R, Guitart-Masip M, Vicens-Costa E, Gimenez-Llort L, Valdar W, Johannesson M, et al. Fearfulness in a large N/Nih genetically heterogeneous rat stock: differential profiles of timidity and defensive flight in males and females. Behav Brain Res. 2008;188:41–55.

75. Pestana JE, Graham BM. The impact of estrous cycle on anxiety-like behaviour during unlearned fear tests in female rats and mice: A systematic review and meta-analysis. Neurosci Biobehav Rev. 2024;164:105789.

76. Chan K, Labruijere S, Garrelds I, Danser A, Villalón C, MaassenVanDenBrink A. Measurements of 17β-estradiol levels in mice for migraine research. The Journal of Headache and Pain. 2013;14:P92.

